# No evidence for associations between brood size, gut microbiome diversity and survival in great tit (*Parus major*) nestlings

**DOI:** 10.1101/2022.09.06.506880

**Authors:** M. Liukkonen, M. Hukkanen, N. Cossin-Sevrin, A. Stier, E. Vesterinen, K. Grond, S. Ruuskanen

## Abstract

**Background:** The gut microbiome forms at an early stage, yet data on the environmental factors influencing the development of wild avian microbiomes is limited. As the gut microbiome is a vital part of organismal health, it is important to understand how it may connect to host performance. The early studies with wild gut microbiome have shown that the rearing environment may be of importance in gut microbiome formation, yet the results vary across taxa, and the effects of specific environmental factors have not been characterized. Here, wild great tit (*Parus major*) broods were manipulated to either reduce or enlarge the original brood soon after hatching. We investigated if brood size was associated with nestling bacterial gut microbiome, and whether gut microbiome diversity predicted survival. Fecal samples were collected at mid-nestling stage and sequenced with the 16S rRNA gene amplicon sequencing, and nestling growth and survival were measured.

**Results:** Gut microbiome diversity showed high variation between individuals, but this variation was not significantly explained by brood size or body mass. Additionally, we did not find a significant effect of brood size on body mass or gut microbiome composition. We also demonstrated that early handling had no impact on nestling performance or gut microbiome. Furthermore, we found no significant association between gut microbiome diversity and short-term (survival to fledging) or mid-term (apparent juvenile) survival.

**Conclusions:** We found no clear association between early-life environment, offspring condition and gut microbiome. This suggests that brood size is not a significantly contributing factor to great tit nestling condition, and that other environmental and genetic factors may be more strongly linked to offspring condition and gut microbiome. Future studies should expand into other early-life environmental factors e.g., diet composition and quality, and parental influences.

## INTRODUCTION

The digestive tract hosts a large community of different microorganisms (i.e., gut microbiome) and is known to be a fundamental part of organismal health and a powerful proximate mechanism affecting host performance [1–2]. The gut microbiome has been studied across a wide range of animal taxa e.g., humans [3–5], fish [6], and economically important species such as poultry [7], and data from wild populations is slowly increasing [8]. Generally, a more diverse gut microbiome is considered beneficial for individual health [9], but there are also community structure effects that define the functionality [10]. For example, laboratory-bred mice with a less diverse gut microbiome have a substantially lower chance of surviving an influenza infection compared to their wild counterparts unless receiving a gut microbiota transplant from their wild counterparts [11–12]. Moreover, gut microbiome had been linked to host fitness and survival in the Seychelles warbler (*Acrocephalus sechellensis*). Individuals that harbored opportunistic pathogens, i.e., microbes that usually do not cause disease in healthy individuals, but may become harmful in individuals that are immunocompromised, in their gut microbiome showed higher mortality [13–14]. Therefore, understanding how gut microbiome affects fitness within and between individuals is necessary for not only understanding species survival but also evolution [15–17].

Gut microbiome forms at a young age and remains somewhat stable in adulthood as found for example in laboratory bred mice [18–20]. Disruption in the gut microbiome that leads to a microbiome imbalance at a young age could result in both short-term and long-term changes in the gut microbiome [21–22]. Of the environmental effects, diet [23], including e.g., macronutrient balance (carbohydrates, fats, amino acids) [3, 24] has been concluded to be major determinants of rat and mouse gut microbiome, and this effect has recently been seen in avian models as well [25–28]. Moreover, macronutrient balance has been linked to intestinal microbiome composition [3, 24] and the functioning of individual immune response [29–30]. However, as a large part of the prior research has focused strictly on humans or species living in controlled environments in which environmental effects on both the microbiome and host are sidelined [31–32], many species, including most birds [8], have only started to attract attention [33].

The mechanisms of bacterial colonization of the bird gut are somewhat unique as avian life-histories differ significantly from those of e.g., mammals [34]. In mammals, the offspring are exposed to bacterial colonization during vaginal birth [35] and lactation [36–37], whereas bird hatchlings are exposed to bacteria first upon hatching [20, 38]. Few studies have investigated the possibility of bacterial colonization *in ovo*, but results are still lacking [39]. Genetics [40–42] as well as the post-hatch environment [20, 43–46] have a significant effect on the formation of the avian gut microbiome. Once hatched, most altricial birds feed their young, which exposes the hatchlings to various bacteria that originate from the parents i.e., via vertical transmission [47]. It has also been shown that environmental factors are major contributors in the formation of gut microbiome [48–51], one of these being the rearing environment in the nest [44].

As early-life environment is connected to the establishment of gut microbiome, brood size may affect gut microbiome [52]. Brood size is often associated with parents’ performance and ability to feed their young [53], and the trade-off between offspring quality and quantity has been studied widely [54–55]. Food quantity per nestling can decrease in enlarged broods, as parents may not be able to fully compensate for the additional amount of food an enlarged brood requires [56–57]. For example, in great tits (*Parus major*) it has been shown that nestlings from reduced broods may have a higher body mass [58] and tend to survive better [59]. Importantly, great tit nestling body mass has been connected to gut microbiome diversity and composition: body mass positively correlates with gut microbiome richness [52]. This could imply that good physiological condition and high food availability would allow the host to have a diverse gut microbiome that promotes a healthy gut.

Alterations in early-life gut microbiome could have long-term consequences on individual performance [60], yet such effects have rarely been studied in wild organisms. In wild birds, some bacterial taxa have been linked to better survival. For example, a high abundance of bacteria in the order *Lactobacillales* of the phylum *Firmicutes* is related to higher individual fitness in Seychelles warblers [14] and great tits [61]. These bacteria are also known for the benefits for bird health in economically important species such as poultry, in which *Lactobacilli* are used as probiotics to boost immune functioning [62]. Besides *Lactobacillales,* gut bacteria belonging to other genera such as *Clostridium* and *Streptococcus* are important for the degradation of non-starch polysaccharides and for the synthesis of essential molecules such as the short-chain fatty acids [63–64]. Short-chain fatty acids are important in host energy metabolism [65] and therefore crucial for performance. Changes in nestling’s early-life gut microbiome could affect such key physiological processes that could influence for example nestling body mass, which is tightly linked to survival to fledging [58–59]. Because the gut microbiome establishes at a young age and is less plastic later in life [18–20], gut microbiome and changes to its richness can have long-term effects on juvenile and adult survival [21–22]. For example, antibiotic treatment at infancy can affect the expression of genes involved in immune system functioning and lead to long-term effects on host metabolism [20]. Moreover, changes in the rearing environment can affect individual physiology and these effects can carry over to later stages of an individual’s life such as survival to fledging and lifetime reproductive success [66].

Here, we use an experimental approach to investigate whether brood size manipulation influenced wild great tit nestlings’ bacterial gut microbiome diversity on day 7 post-hatch. We also investigated whether brood size influenced nestling body mass on day 7 or on day 14 post-hatch, and if the gut microbiome predicts short-term (i.e., survival to fledging) and mid-term (i.e., apparent juvenile) survival. The great tit is a well-studied species in the fields of ecology and evolution, and it is easy to monitor in the wild due to its habit of breeding in nest boxes. Great tit nestlings’ gut microbiome undergoes profound shifts during early life [52], and it has been linked to nestling natal body mass and body size [52, 61], yet studies focusing on gut microbiome associations with survival are still scarce. Here, we manipulated wild great tit broods by reducing or enlarging the original brood size in order to analyze if this affected the gut microbiome. In large broods, nestlings need to compete for their food more [67–68], and the lower food availability could result in a lower gut microbiome diversity. This might impair nestling body mass and fitness prospects [13, 52]. We used a partial cross-fostering design that enabled us to disentangle the relative contributions of genetic background, early maternal effects, and rearing environment such as parents, nest and nestmates on gut microbiome. Furthermore, we used an unmanipulated control group in which no nestling was cross-fostered to control for the possible effects of moving the nestlings between nests. For example, early human handling such as marking and weighing at day 2 post-hatch could influence gut microbiome later on. We hypothesized that 1) in reduced broods nestlings would have a higher body mass, 2) in reduced broods nestling gut microbiome would be more diverse than in enlarged broods, and 3)higher gut microbiome diversity on day 7 post-hatch would increase survival to fledging and potentially reflect apparent juvenile survival. Such knowledge could provide new information about gut microbiome in wild passerine bird population and how the early-life environment may associate with nestling gut microbiome, body mass, and short-term and mid-term survival.

## METHODS

### Study area and species

The great tit is a small passerine bird, which breeds in secondary holes and artificial nest-boxes, making it a suitable model species. Great tits breed throughout Europe and inhabit parts of Northern Africa and Asia as well, and the breeding areas differ in environment and diet [69]. In Finland the great tit is a common species with an estimate of 1.5 to 2 million breeding pairs. They lay 6 to 12 eggs between April and May and the female incubates the eggs for 12 – 15 days. The nestlings fledge approximately 16 to 21 days after hatching. The study was conducted during the breeding season (May-July 2020) on Ruissalo island (60° 25’ 59.99" N 22° 09’ 60.00" E). Ruissalo island habitat is a mostly temperate deciduous forest and meadows, and some areas have small patches of coniferous trees.

### Brood size manipulation experiment

Nest boxes were first monitored weekly and later daily when clutches were close to the estimated hatching date. Brood size manipulation took place on day 2 after hatching. Increases in great tit brood size can lead to lowered weight in both the nestlings and adults [70–75], and our decision on the number (i.e., +2 or -2) of manipulated nestlings (i.e,. +2 or -2) followed the cited studies. We had four treatment groups (see Fig. 1): in the ‘enlarged group (henceforward called E)’, we increased the brood size by two individuals that were taken from a ‘reduced brood’. Correspondingly, in the ‘reduced group (henceforward called R)’, we decreased the brood size by two individuals, that were added to the enlarged broods. In the ‘control group (henceforward called C)’, we swapped nestlings between nests but did not change the brood size. And lastly, in the ‘unmanipulated control group (henceforward called COU)’, we only weighed and collected fecal samples on day 7 but did not move the nestlings between nests. We also moved nestlings between the reduced nests to ensure that all nests except for COU had both original and fostered nestlings. Control nests were used to control for potential cross-fostering effects unrelated to brood size. Additionally, in the unmanipulated control group nestlings were not moved or weighed on day 2 in order to control for any handling effects *per se*. This study design enabled us to test the potential impacts of handling nestlings and swapping the nest early after hatching. We aimed to move approximately half of the chicks in the manipulated nests, so that the number of original and the fostered nestlings would be the same in each nest after manipulation.

**Figure 1.**
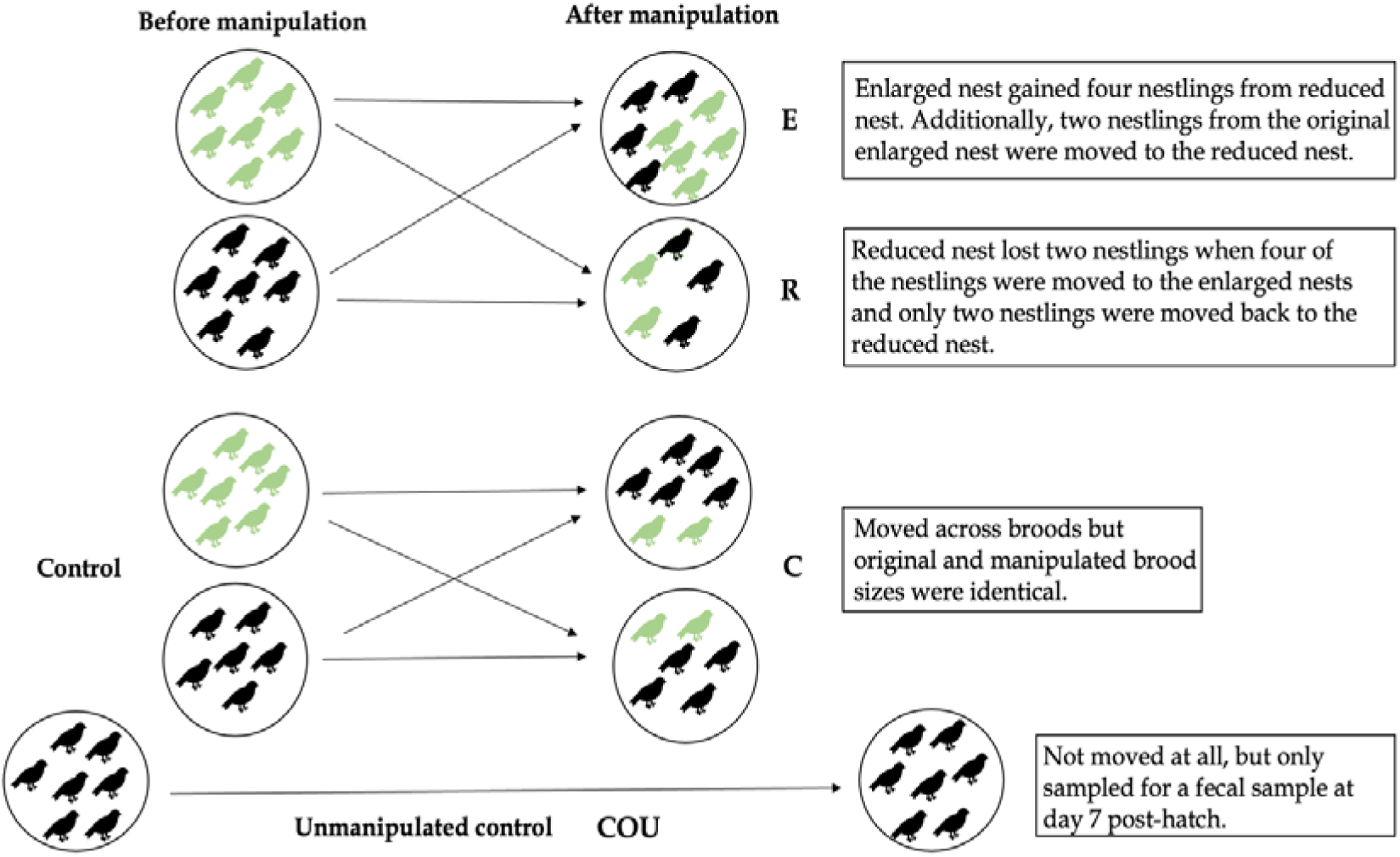
Brood size manipulation experiment schematic diagram. 2-day-old nestlings were moved between nestboxes to enlarge or to reduce original brood size (an example with brood size of seven is given). Some nests were kept as control nests (nestlings were moved but brood size remained the same) and some were kept as unmanipulated control nests (nestlings were not moved at all to test whether early-life handling affects gut microbiome). The original brood size varied between nests.

Before they were moved, nestlings were weighed using a digital scale with a precision of 0.1 g and identified by clipping selected toenails. We aimed to add/remove nestlings that were of similar weight to avoid changing the sibling hierarchy in the brood. The moving procedure was performed as quickly as possible to minimize the risk of stress and the nestlings were kept in a warmed box during transportation. For each pair of nests in the brood size manipulation experiment, we selected nests that had a similar hatching date. In case of uneven number of nests hatching within a day, one or three nest(s) was/were allocated to the COU group. To avoid potential bias from hatching date, we allocated nests in any given day evenly to each treatment. We also checked that the treatments had an equal brood size on average i.e., we did not want to only reduce the larger clutches and enlarge the smaller clutches. These is also a significant bias towards COU nests being later in the season on average (Table 1).

**Table 1.**
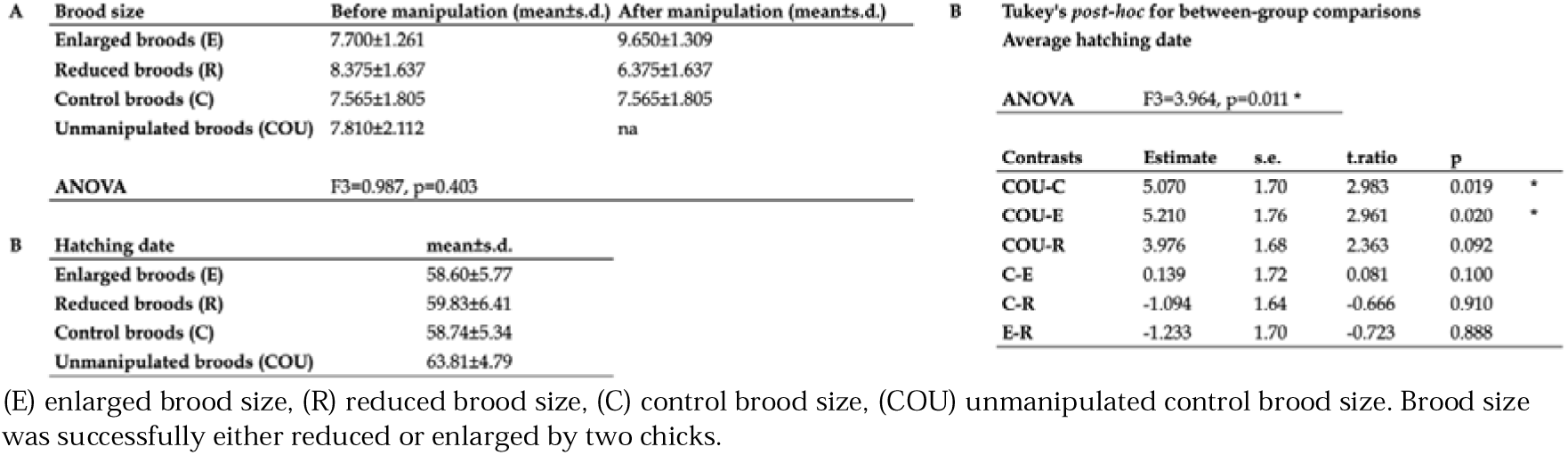
(A) Brood size before and after manipulation, (B) Hatching date across treatments.

### Fecal sample collection

To study the effects that brood size may have on the nestling gut microbiome and its links to individual nestling body mass, survival to fledging and apparent juvenile survival, we used a subset of data from a larger experiment (Cossin-Sevrin et al., *unpublished data*). In this subset, we use individuals from which fecal samples were collected on day 7 after hatching and analyzed for microbiome diversity and composition (C=23 nestlings/15 nests, COU=22/13, E=23/15, R=24/16) We aimed to collect two samples (one from original and one from foster nestlings) per nest. Fecal samples from the nestlings were collected gently by stimulating the cloaca with the collection tube. Samples were collected straight into a sterile 1.5 ml Eppendorf tube to avoid possible contamination of the sample. At time of sampling, each nestling was weighed (0.1g), and the nestlings were ringed for individual identification using aluminum bands. The samples were stored in cool bags onsite and afterwards moved to a -80 °C freezer for storage until DNA extraction.

### Apparent juvenile survival

We monitored all study nests until fledging to measure short-term survival. On day 14 post-hatch, the sampled nestlings were weighed, and wing-length was measured to detect if the manipulation had any effects on nestling growth. Nests were subsequently monitored for fledging success. Additionally, we monitored our study population for apparent juvenile survival (i.e., mid-term survival) after the breeding season (i.e., approximately 3 months after fledging) to assess the association between gut microbiome and post-fledging survival. We captured juvenile great tits by mist netting during the autumn-winter 2020 at six different feeding stations that had a continuous supply of sunflower seeds and suet blocks. Feeding stations were located within the previously mentioned nest box population areas. For each site mist netting with playback was conducted on three separate days during October-November 2020 for three hours at a time, leading to a total of 69 hours of mist netting. A total of 88 individuals from the brood size manipulation experiment were caught, and the caught juvenile great tits were weighed, and wing length was measured. Our catching method provides an estimate of post-fledging survival yet, it could be slightly biased based by dispersal. In a previous study in our population [76], none of the birds ringed as nestlings were recaptured outside the study area, suggesting that dispersal is likely limited.

### DNA extraction and sequencing

We chose two samples per nest for DNA extraction, yet in such a way that both fledged and not-fledged nestlings would be included in the dataset. DNA was extracted from nestling fecal samples using the Qiagen QIAamp PowerFecal Pro DNA Kit (Qiagen; Germany) following the manufacturer’s protocols. Additionally, we included negative (RNAse and DNAse free ddH_2_O) controls to control for contamination during DNA extraction and additional controls to confirm successful amplification during PCR. A short fragment of hypervariable V4 region in the 16S rRNA gene was amplified using the purified DNA samples as template with the following primers: 515F_Parada (5’ - GTGYCAGCMGCCGCGGTAA -3’) and 806R_Apprill (5’ - GGACTACNVGGGTWTCTAAT - 3’) [77–78]. PCRs were performed in a total volume of 12 µL using MyTaq RedMix DNA polymerase (Meridian Bioscience; Cincinnati, OH, USA). The PCR cycling conditions were as follows: first, an initial denaturation at 95 °C for 3 minutes followed by 30 cycles of 95 °C for 45 seconds, 55 °C for 60 seconds, and 72 °C for 90 seconds, and finished with a 10-minute extension at 72 °C. After the first round of PCR, a second round was conducted to apply barcodes for sample identification [79]. For this, PCR cycling conditions were as follows: first, an initial denaturation at 95 °C for 4 minutes followed by 18 cycles of 98 °C for 20 seconds, 60 °C for 15 seconds, and 72 °C for 30 seconds, and finished with a 3-minute extension at 72 °C. We performed replicate PCR reactions to control for errors during the amplification. Further on, the PCR products were measured for DNA concentration with Quant-IT PicoGreen dsDNA Assay Kit (ThermoFischer Scientific; Waltham, MA, USA) and for quality with TapeStation 4200 (Agilent; Santa Clara, CA, USA). The samples from each of the PCR replicates were pooled equimolarly creating two separate pools and purified using NucleoMag NGS Clean-up and Size Select beads (Macherey-Nagel; Düren, Germany). Finally, pooled samples were sequenced (2 x 300 bp) on the Illumina MiSeq platform (San Diego, CA, USA) at the Finnish Functional Genomic Center at the University of Turku (Turku, Finland).

### Sequence processing

All statistical analyses were performed with R (v. 4.11.0; R Development Core Team 2021) unless otherwise stated. The demultiplexed Illumina sequence data was first processed with Cutadapt version 2.7 [80] to remove locus-specific primers from both R1 and R2 reads. Then, the DADA2 pipeline (v. 1.24.0; [81]) was used to filter the reads based on quality, merge the paired-end (R1 and R2) reads, to define the single DNA sequences i.e., Amplicon Sequence Variants (henceforward ASV), and to construct a ‘seqtab’. Seqtab is a matrix also known as otutable or readtable: ASVs in columns, samples in rows, number of reads in each cell, using default parameter settings. In total, our seqtab consisted of 6,929,537 high-quality reads. Reads were assigned to taxa against the SILVA v132 reference database [82] resulting in 8658 ASVs. To control for contamination, negative DNA extraction and PCR controls were used to identify contaminants (60 ASVs) using the decontam package (v. 1.12; [83]) and all were removed from the dataset. Sequencing runs (replicate PCR’s) were merged using the phyloseq package (v. 1.32.0) and non-bacterial sequences (mainly *Chlorophyta*) were removed from the data as they were not of interest in this study resulting in a total of 6566 ASVs (a total of 4,107,028 high-quality reads in all samples; mean per sample: 15,155.085; mean range per sample: 0-97,264). Singleton reads were removed from the dataset by the DADA2 pipeline. Data was further analyzed with the phyloseq package (v. 1.32.0; [84]), and the microbiome package (v. 1.18.0; [85]) and visualized with the ggplot2 package (v. 3.3.6; [86]).

The final dataset contained 92 samples from great tit nestlings resulting in a total of 3,161,696 reads (mean per sample: 34,366.261; mean range per sample 108 – 189,300 reads), which belonged to 6,505 ASVs. The dataset was then rarefied for alpha diversity analyses at a depth of 5000, as this was where the rarefaction curves plateaued (see Supplementary file 2). The rarefied dataset contained 4,791 ASVs in 88 samples. For beta diversity, the unrarefied dataset was used after confirming that the beta diversity statistics were quantitatively similar for the rarefied and unrarefied datasets. Bacterial relative abundances were summarized at the phylum and genus level and plotted based on relative abundance for all phyla and genera. A Newick format phylogenetic tree with the UPGMA algorithm to cluster treatment groups together was used to visualize sample relatedness (see Supplementary file 3) and was constructed using the DECIPHER (v. 2.24.0; [87]), phangorn (v. 2.8.1; [88]), and visualized with ape (v. 5.6-2; [89]), and ggtree (v. 3.4.0; [90]) packages.

## Statistical analyses

### Nestling body mass

First, to analyze whether brood size manipulation affected nestling body mass in the C, E, and R treatment groups, we ran two linear mixed-effects models with the *lme4* package (v. 1.1-29; [91]). In these models we used either body mass on day 7 or 14 as the dependent variable and brood size manipulation treatment, hatching date, body mass on day 2 post-hatch and original brood size as predicting variables. Hatching date is used as a predicting variable because it is known to affect nestling body mass during the breeding season [92] and there were differences in hatching date between the COU and other treatment groups (see Table 1). We included the interaction between original brood size and brood size manipulation treatment in both models as the effect of manipulation may depend on the original brood size. For example, there could be stronger effect of enlargement in already large broods. Nest of origin and nest of rearing were used as random intercepts to control for the non- independence of nestlings sharing the same original or foster nests. Here, we did not include the COU group in the analysis because we wanted to measure the effects of treatment on nestling body mass, and only enlarged, reduced or control broods’ nestlings were moved between nests.

Second, to analyze whether the actual brood size affected nestling body mass, we ran two models where we used it as a continuous dependent variable to explain body mass either on day 7 or on day 14 post-hatch. Hatching date and body mass on day 2 post-hatch were used as predicting variables and nest of origin and nest of rearing as random intercepts to control for the non-independency of samples. We included the interaction between manipulated brood size and hatching date in the models because the effect of brood size may depend on the hatching date. For example, hatching date can reflect environmental conditions and large broods may perform poorly late in the season due to poorer food availability. The COU group was initially excluded from this model to see which of the two random effects, nest of origin or nest of rearing, explained a larger portion of variation in the treatment groups. In the COU group, nest of origin and nest or rearing were the same, which meant we could not include both random effects in models where all treatment groups were present due to the model failing to converge. Nest of origin explained more of the variation in the first model (see Supplementary file 4) and therefore, we used it in the full models with all treatment groups: C, COU, E and R. In these models, nestling body mass either on day 7 and or on day 14 post-hatch was used as a dependent variable and manipulated brood size as the explanatory variable. Hatching date and body mass on day 2 post-hatch were set as predicting variables. Nest of rearing was used as a random intercept to control for the non-independence of nestlings sharing the same foster nests. The significance of factors included in the models were tested using the F-test ratios in analysis of variance (Type III ANOVA).

### Alpha diversity

For alpha diversity analyses, which measures within-sample species diversity, we ran two linear mixed-effects models with the *lme4* package (v. 1.1-29; [91]) to measure if either brood size manipulation or manipulated brood size as a continuous variable were associated with gut microbiome diversity. We used two alpha diversity metrics: the Shannon Diversity Index, which measures the number of bacterial ASVs and their abundance evenness within a sample, and Chao1 Richness, which is an estimation of the number of different bacterial ASVs in a sample. Both metrics were used to check if alpha diversity results were consistent across different metrics. Each diversity index was used as the dependent variable at a time and either brood size manipulation treatment or manipulated brood size as a predicting variable. In both models we included original brood size, weight on day 7 post-hatch and hatching date as covariates. We included interaction between brood size manipulation treatment and original brood size as there could be a stronger effect of enlargement in initially large broods. We also included interaction between manipulated brood size and weight on day 7 post-hatch because effect of brood size on microbiome may depend on nestling weight. We also tested whether alpha diversity predicted weight on day 7 post-hatch, as weight and gut microbiome diversity have been connected in previous studies. In this analysis we used weight on day 7 post-hatch as the dependent variable and alpha diversity (Shannon Diversity Index and Chao1 Richness), treatment and hatching date as predicting variables and nest of rearing as the random effect. In these sets of models, we first excluded the COU group to see which of the two random effects, nest of origin or nest of rearing, explained a larger proportion of variation in the treatment groups. Nest of rearing explained more of the variation in this model (see Supplementary file 4) and therefore, we used it in the full model with all treatment groups: C, COU, E and R. The significance of factors included in the models were tested using the F-test ratios in analysis of variance (ANOVA).

### Short-term survival

To explore whether alpha diversity associated with survival to fledging (i.e., short-term survival) and with apparent juvenile survival in Autumn 2020 (i.e., mid-term survival), we used generalized linear models with binomial model (v. 1.1-29; lme4 package, [91]), and then tested the significance of factors with type 2 ANOVA from the car package (v. 3.0-13; [93]). Type 2 ANOVA was used because the model did not contain interaction between predicting and there was no order between covariates, as they could not be ranked. Survival to fledging and recapture in Autumn 2020 were used as the binomial response variable (yes-no) in each model. Alpha diversity (Shannon Diversity Index and Chao1 Richness) was the main predicting variable, and weight on day 7 post-hatch (same time as sampling the fecal gut microbiome), hatching date and manipulated brood size were included as covariates in the model. We did not include brood size manipulation treatment in the survival models as not enough birds from each treatment group were recorded for fledging and juvenile survival. Moreover, we excluded random effects from this model as the model failed to converge. 65 nestlings fledged successfully, while 8 nestlings were found dead in nest boxes. For 15 nestlings we had no fledging record, so these were excluded from the survival to fledging analysis. In apparent juvenile survival, 19 birds out of 92 (with data on microbiome diversity) were recaptured as juveniles. For all analyses, the R package car (v. 3.0-13; [93]) was used to test Variance Inflation Factors (VIFs) and the package DHARMa (v. 0.4.5; [94]) to test model diagnostics for linear mixed-effects and generalized linear models.

### Beta diversity

For visualizing beta diversity, i.e., the similarity or dissimilarity between the treatment group gut microbiomes, non-metric multidimensional scaling (NMDS) was used with three distance matrices: Bray-Curtis [95], weighted UniFrac, and unweighted UniFrac [96]. Permutational multivariate analysis of variance (PERMANOVA) using the Euclidean distance matrix and 9999 permutations was tested with the R package vegan (*adonis2* function; v. 2.6-2; [97]) to investigate if any variables affected to the variation in gut microbiome composition. Nest of rearing was set as a blocking factor in the PERMANOVA to control for the non-desirable effects of the repeated sampling of foster siblings. The test for homogeneity of multivariate dispersions was used to measure the homogeneity of group dispersion values. We used the phyloseq package (v. 1.32.0; [84]) to run a differential abundance analysis with a significance cut-off p < 0.01 to test the differential abundance of ASVs between the treatment groups.

## RESULTS

### The effects of brood size manipulation on nestling body mass

Brood size manipulation did not significantly affect nestling body mass on day 7 post-hatch (ANOVA: F_2, 25.832_=0.441, p=0.648; see Supplementary file 5). Moreover, there was no significant interaction between brood size manipulation and original brood size (ANOVA: F_2, 24.610_=0.678, p=0.517; see Supplementary file 5). On day 14 post-hatch, brood size manipulation did not significantly affect nestling body mass (ANOVA: F_2, 24.335_=0.831, p=0.448; see Supplementary file 5). However, body mass increased with increasing hatching date (ANOVA: F_1 24.070_=13.367, p=0.001; see Supplementary file 5). Next, we did not find any significant associations between manipulated brood size and nestling body mass (ANOVA for weight on day 7: F_1, 35.149_=1.777, p=0.191; ANOVA for weight on day 14: F_1, 29.491_=2.156, p=0.153; see Supplementary file 6). Nest of origin explained a larger proportion of the variation in weight than the nest of rearing on both day 7 (nest of origin 41.1 % and nest of rearing 24.4 %) and day 14 (nest of origin 65.5 % and nest of rearing 21.9 %) post-hatch, but this result was not statistically significant (Pr > χ2 = 1) (see Supplementary file 4).

### Alpha diversity

As 7-day-old nestlings, most bacterial taxa belonged to the phyla *Proteobacteria*, *Firmicutes*, and *Actinobacteria* (Fig. 2).

**Figure 2.**
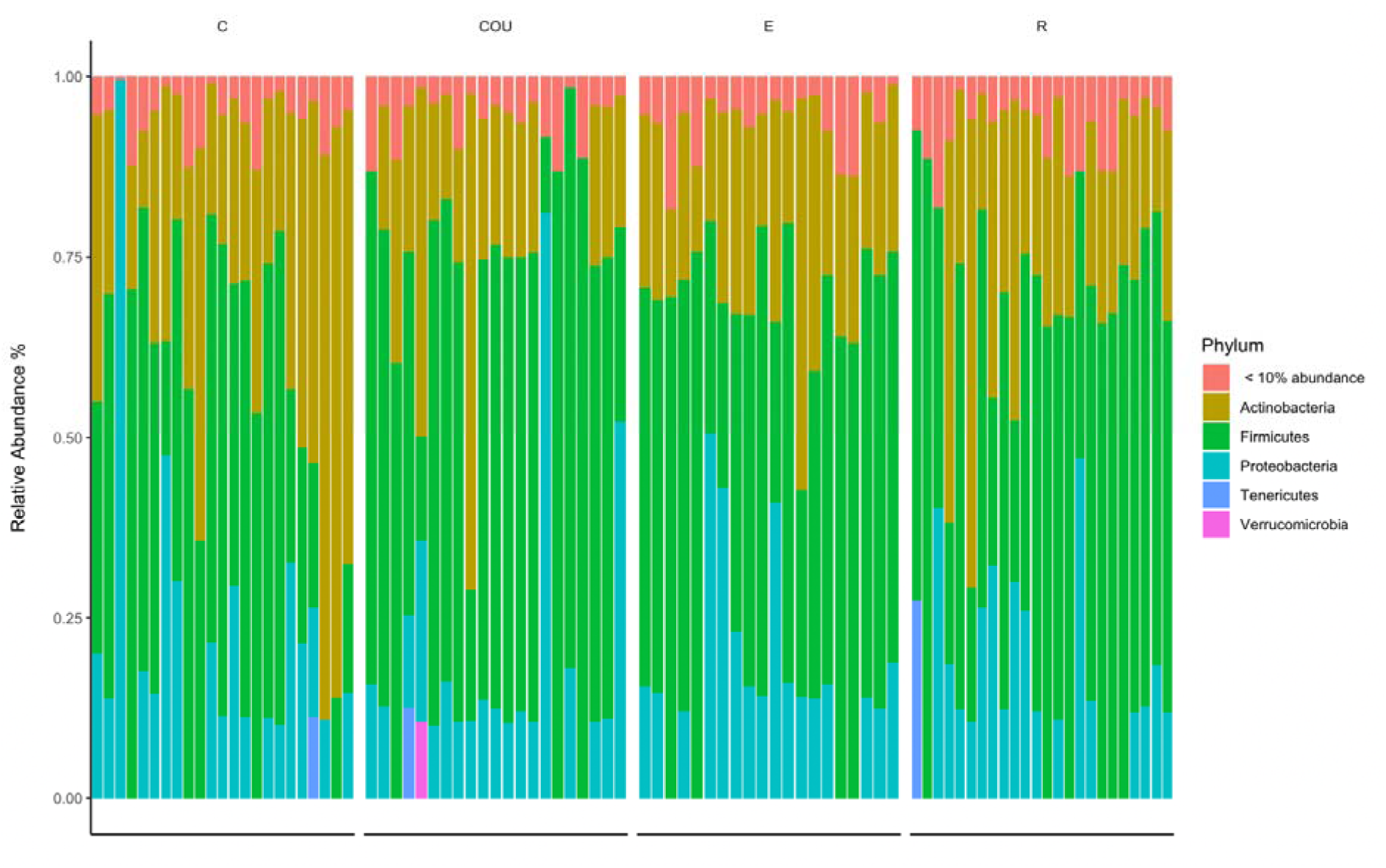
Bacterial relative abundances on Phylum level across the four treatment groups. Each bar represents an individual sample. Treatment groups are control (C), unmanipulated control (COU), enlarged (E), and reduced (R). N=88 samples divided into treatment groups as follows: C=23, COU=21, E=20, R=24. Phyla with less that 10 % in relative abundance is collapsed into the category “< 10 % abundance.”

Brood size manipulation did not significantly influence alpha diversity (Shannon Diversity Index) (ANOVA: F_3, 47.488_=1.026, p=0.390, Fig. 3; see Supplementary file 7). Moreover, original brood size (ANOVA: F_1, 50.269_=0.388, p=0.536; see Supplementary file 7), weight on day 7 post-hatch (ANOVA: F_1, 80.551_=0.003, p=0.959; see Supplementary file 7), and hatching date (ANOVA: F_1, 50.276_=1.073, p=0.305; see Supplementary file 7) did not significantly associate with alpha diversity. There was no significant interaction between brood size manipulation and original brood size (ANOVA: F_3, 48.053_=0.126, p=0.944; see Supplementary file 7). Results for Chao1 Richness were quantitatively similar: brood size manipulation did not affect alpha diversity (ANOVA: F_3, 45.936_=0.358 p=0.784, Fig. 3; see Supplementary file 7). Nest of rearing explained a larger proportion of the observed variance in alpha diversity (27.7 %) than nest of origin (10.8 %), but the result was not statistically significant (Pr > χ2 = 1) (see Supplementary file 4).

**Figure 3.**
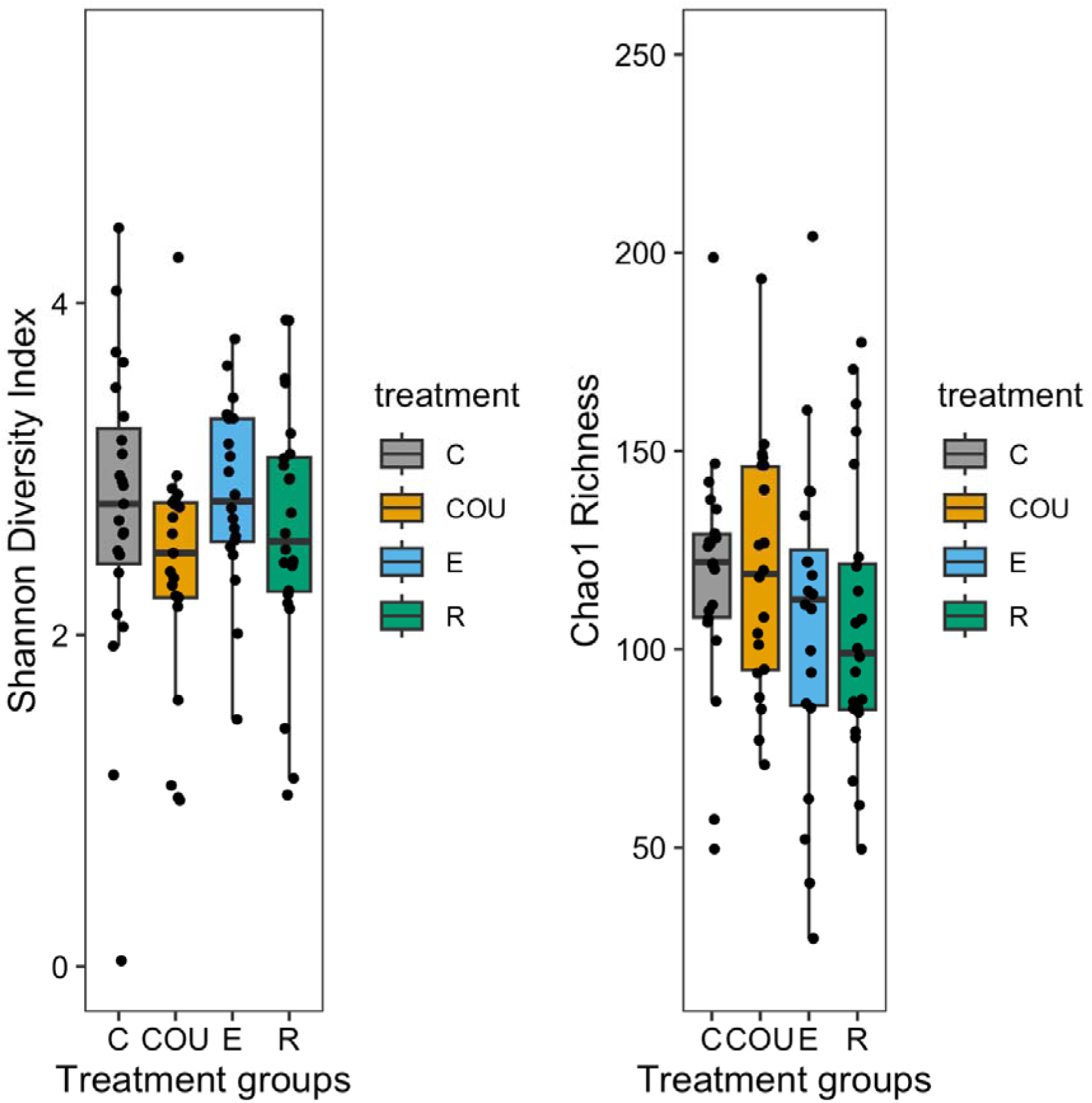
The gut microbiome alpha diversity of 7-day-old great tit nestlings across the four treatment groups visualized with two diversity metrics: A) Shannon Diversity Index and B) Chao1 Richness. The black dots represent each observation within a treatment group. The whiskers represent 95 % confidence intervals. Treatment groups are control (C), unmanipulated control (COU), enlarged (E), and reduced (R). N=88 samples divided into treatment groups as follows: C=23, COU=21, E=20, R=24.

Next, we tested whether the manipulated brood size as a continuous variable was associated with alpha diversity (Shannon Diversity Index), but found no significant association (ANOVA: F_1, 63.001_<0.001, p=0.984; see Supplementary file 8) in this analysis either. Weight on day 7 post-hatch (ANOVA: F_1, 82.840_=0.015, p=0.903; see Supplementary file 8) and hatching date (ANOVA: F_1, 59.734_=0.137, p=0.713; see Supplementary file 8) did not correlate with alpha diversity in this model either. There was no significant interaction between manipulated brood size and weight on day 7 post-hatch (ANOVA: F_1, 82.702_<0.000, p=0.998; see Supplementary file 8). Results for Chao1 Richness were quantitatively similar (ANOVA: F_1, 65.064_=0.246, p=0.622; see Supplementary file 8): manipulated brood size did not affect alpha diversity, and neither did weight on day 7 post-hatch (ANOVA: F_1 83.513_=0.690, p=0.409; see Supplementary file 8) nor hatching date (ANOVA: F_1 57.110_=1.133, p=0.292; see Supplementary file 8).

### Alpha diversity and short/mid-term survival

Next, we explored whether alpha diversity (Shannon Diversity Index and Chao1 Richness) contributed to predicting short/mid-term survival (survival to fledging and apparent juvenile survival). Survival to fledging was not predicted by alpha diversity (Shannon Diversity Index: χ2=0.010, df=1, p=0.923; see Supplementary files 9 and 10), manipulated brood size (χ2=0.090, df=1, p=0.764; see Supplementary file 9), weight on day 7 post-hatch (χ2=0.388, df=1, p=0.533; see Supplementary file 9) or hatching date (χ2=0.438, df=1, p=0.508; see Supplementary file 9).

Apparent juvenile survival was not significantly associated with alpha diversity (Shannon Diversity Index: χ2=1.916, df=1, p=0.166; see Supplementary files 9 and 10). Moreover, there was no significant interaction between alpha diversity and manipulated brood size (χ2= 1.268, df=1, p=0.260; see Supplementary file 9). However, apparent juvenile survival was negatively associated with hatching date (χ2=4.654, df=1, p=0.031; see Supplementary file 9). Additional analyses to check for the consistency of results were tested the following way: survival to fledging with nestlings from the COU group removed and apparent juvenile survival without the nestlings with no recorded survival for fledging (see methods). These results were quantitatively similar as in the whole dataset for both Shannon Diversity Index (survival to fledging: χ2= 2.285, df=1, p= 0.131; apparent juvenile survival: χ2=1.515, df=1, p=0.218; see Supplementary file 11) and Chao1 Richness (survival to fledging: χ2=0.665, df=1, p=0.415; apparent juvenile survival: χ2=2.654, df=1, p=0.103; see Supplementary file 11).

### Beta diversity

Non-metric multidimensional scaling (NMDS) using weighted and unweighted UniFrac and Bray-Curtis dissimilarity did not show clear clustering of samples based on brood size manipulation treatment (Fig. 5; see Supplementary file 3). The test for homogeneity of multivariate dispersions supported the visual assessment of the NMDS (Betadispersion_9999 permutations_: F_3, 0.069_= 0.650, p<0.001; see Supplementary file 12). Pairwise PERMANOVA further indicated that the treatment (PERMANOVA: R^2^=0.061, F=1.951, p=0.278; see Supplementary file 12), weight on day 7 post-hatch (PERMANOVA: R^2^=0.015, F=1.387, p=0.091) or hatching date (PERMANOVA: R^2^=0.0232, F=2.214, p=0.993) did not significantly contribute to the variation in gut microbiome composition between the treatment groups. Differential analysis of ASV abundance between the treatment groups showed that there is variation in taxa abundance. E group showed higher taxa abundance when compared to COU and C groups and was slightly higher than the R group. C and COU groups were generally lower in taxa abundance than R and E groups, and COU group showed lower abundance than the other groups in each comparison (see Supplementary file 13).

## DISCUSSION

In this study, we investigated the associations between great tit nestling gut microbiome, brood size, and nestling body mass by experimentally manipulating wild great tit broods to either reduce or enlarge the original brood size. The results show that even though there was individual variation in the nestling gut microbiome (Fig. 2), brood size did not significantly contribute to gut microbiome diversity. Neither did gut microbiome diversity explain short-term (survival to nestling) nor mid-term (apparent juvenile) survival. Body mass was also not significantly affected by brood size manipulation. The COU group that functioned as a control for moving and handling effects, did not differ in this respect from the other groups. This suggests that human contact or handling nestlings 2 days post-hatch did not influence nestling gut microbiome or body mass. The partial cross-fostering design enabled us to disentangle the relative contributions of rearing environment (i.e., parents, nest and nestmates) from genetic, prenatal such as maternal allocation to egg, and early post-natal effects such as feeding up to day 2. Nest of rearing seemed to explain more of the variation in nestling gut microbiome diversity than the nest of origin (although not statistically significant), which follows previous studies. Contrastingly, nest of origin seemed to be a stronger contributor than the nest of rearing on nestling body mass on day 7 and day 14 post-hatch. This result was also not statistically significant.

### Brood size manipulation and nestling body mass

First, we explored whether brood size was associated with nestling body mass, as such changes may explain the underlying patterns in gut microbiome [52]. Against our hypothesis, we found no significant association between nestling body mass and brood size: neither reduction nor enlargement of the broods resulted in significant body mass differences in the nestlings on day 7 and day 14 post-hatch. While the result is supported by some studies in which associations between nestling body mass and brood size have been tested [61, 98], the majority of the literature shows that brood size negatively correlates with nestling body mass: in larger broods nestlings are generally of lower mass [52-53, 57, 67, 99-104].

There are a few possible explanations why brood size manipulation did not affect nestling body mass. Firstly, if environmental conditions were good, parents may have been able to provide enough food even for the enlarged nests and thus, variance in brood size may not result in differences in nestling body mass between reduced and enlarged nests. In that case the number of nestlings transferred between enlarged and reduced nests should probably have been larger to create differences in nestling body mass between the two treatments. Still, we think that the decision to transfer +2/-2 was reasonable since it was based on extensive evidence from previous studies [103]. Secondly, it could be that the enlarged brood size negatively influences some other physiological traits while body mass was retained at the expense of these other traits e.g., immune system functioning [105–106]. Moreover, our analysis showed that hatching date had a significant effect on nestling body mass: nestlings that hatched later in the season were of lower weight. This could be a result of changes in the food items that great tits use, changes in temperature conditions or in parental investment during the breeding season. As the season progresses, the abundance of insect taxa varies, and this can result in changes in nutrient rich food [103, 107]. For example, great tits can select certain lepidopteran larvae that vary in their abundance during the great tit breeding season [108]. Thirdly, it could be that the change in brood size was influencing the parents’ condition instead of the nestlings [109–110]. In enlarged broods, parents are required to forage more which can lead to higher energy expenditure and increased stress levels in parents [72–73, 109].

### Brood size manipulation and gut microbiome

We found large inter-individual differences in gut microbiome diversity, yet this variation was not explained by brood size or nestling body mass. It is possible that brood size did not result in differences in food intake. For example, parents were likely able to provide an equivalent amount of food, given that body mass was not significantly affected by the brood size manipulation. Therefore, brood size manipulation did not affect gut microbiome diversity through differences in nutrient uptake. Alternatively, in this study, fecal sampling took place 5 days after the initial brood size manipulation (day 2 post-hatch). It could be that sampling on a later date or at multiple timepoints [61, 111] would have led to different results. Firstly, the time interval may not have been long enough to detect effects of the brood size manipulation. Secondly, it has been shown in previous studies that the nestling gut microbiome undergoes profound shifts at the nestling stage: overall gut microbiome diversity decreases but relative abundance in some taxa increases [52]. We suggest that fecal samples could be collected on multiple days post-hatch to understand the potential day to day changes in the nestling gut microbiome.

Our results suggest that the variance in gut microbiome is a result of other factors than those linked to brood size. Firstly, one of these factors could be diet (i.e., food quality) which has gained attention in gut microbiome studies during the past years [25, 27, 112–115]. The overall diversity in gut microbiome could be explained by adaptive phenotypic plasticity because it is sensitive to changes in the environment e.g., changes in diet [116–117]. The food provided by the parents can vary between broods in different environments [118], and this variation in diet can lead to differences in gut microbiome diversity [114–119]. For example, abundance in certain dietary items such as insects or larvae can result in lower gut microbiome diversity than other dietary items [113–116]. As great tits have been reported to adapt their diet along the breeding season due to changes in insect taxa frequency [103, 107] this could affect the between-nestling and between-nest gut microbiome diversity. However, using wild bird populations in gut microbiome studies limits the ability to control the consumed dietary items because parents may use variable food resources and there can be variance in dietary between sexes and even individuals. Visual assessment of dietary items [116] and metabarcoding could be of use here as they enable the identification of food items on genus and even species level from e.g., fecal samples [119].

Secondly, breeding habitat may lead to differences in gut microbiome diversity [120]: adult birds living in deciduous forests have shown to harbor different gut microbiome diversity than their counterparts living in open forested hay meadows. Here, we used a cross-fostering design to study if the rearing environment contributed to the variation in gut microbiome diversity: Our study indicated that the nest of rearing seemed to explain more of the gut microbiome variation than the nest of origin (although not significant), which follows some previous results [43–44, 52]. For example, a study with great and blue tit (*Cyanistes caerulaeus*) nestlings showed that the nest of rearing contributed more to the gut microbiome than the nest of origin [43], and another study with the brown-headed cowbird (*Molothrus ater*) concluded that the sampling locality had a significant contribution to the gut microbiome [44]. Teyssier et al. [52] conducted cross-fostering at day 8 post-hatch in great tits and found that the nest of rearing influenced the gut microbiome more than the nest of origin. Additionally, parents can pass down their bill and feather microbiome through vertical transmission, which could influence nestling gut microbiome [20].

Results from beta diversity analysis were similar to that of alpha diversity: brood size manipulation did not contribute to the variation in gut microbiome composition. Overall, variation in gut microbiome composition could be a result of different genetic and environmental contributors. Firstly, great tit nestling gut microbiome composition could be explained by underlying genetic effects that we did not measure in this study. Phylosymbiosis i.e., the matching of gut microbiome composition to host genetic structure, could be explained by underlying genetics that may translate into physiological differences that affect the gut microbiome e.g., founder effects or genetic drift [121]. Davies et al. [14] found that MHC genes correlate with gut microbiome composition: the expression of specific alleles in the MHC genes was connected to the abundance of specific bacterial taxa such as *Lactobacillales* and *Bacteroidales* that influenced host health. In a study by Benskin et al. [41] captive zebra finches (*Taeniopygia guttata*) showed significant variation in gut microbiome composition between individuals even though their diet and housing conditions were standardized. The study suggested that individual homeostatic mechanisms linking to naturally occurring differences in individual gut microbiome could be why gut microbiome composition varied even with standardized housing conditions [41]. Secondly, gut microbiome composition could have been affected by the same environmental effects that may have linked to the variation in gut microbiome diversity: diet and feeding behavior [115–116].

Differential analysis of ASV abundance showed variation in differential abundance of taxa between the treatment groups. However, several ASVs were not taxonomically assigned beyond family level making it difficult to draw conclusions about the significance of these results. All treatment groups had taxa belonging to the order *Firmicutes, Proteobacteria* and *Actinobacteria*, which was to be expected because they are usually the most core phyla in passerine gut microbiomes [33]. Nestlings belonging to E, R or C group showed higher taxa abundance than the COU group in each comparison. This result could be a result of the COU nestlings generally hatching later in the season and potentially having a less diverse diet [103, 107]. Of the E, R and C groups, C group was less abundant than E and R groups. Both E and R group showed high taxa abundance, which is interesting because we hypothesized that nestlings belonging to the E group would potentially experience less parental investment per nestling and have lower gut microbiome diversity and therefore, be less abundant [56-57, 67-68]. We did not observe differential abundance in e.g., the order *Lactobacillales* which would have been of interest, because the order hosts taxa that are beneficial for gut microbiome health [14, 62]. The genus *Staphylococcus* was differentially abundant in the E group, but not in the other groups. *Staphylococcus* is a gram-positive genus of bacteria and known to cause infections in its host species [122]. Curiously, the COU group was differentially abundant in the genus *Dietzia*, which is a human pathogen [123].

### Gut microbiome and short-term and mid-term survival

Our results showed that gut microbiome diversity and brood size were not significantly associated with short-term (survival to fledging) or mid-term (apparent juvenile) survival. However, while a more diverse gut microbiome is considered a possible indicator of a healthy gut microbiome, the effects of the gut microbiome on the host health may often be more complex and related to specific taxa [9–10]. For example, Worsley et al. [13] did not find a correlation between body condition and gut microbiome diversity, yet they found that specific taxa in the gut microbiome linked with individual body condition and survival. Not only environment, but also genetic background of the individual may contribute to gut microbiome and survival. In a study by Davies et al. [14], *Ase-ua4* allele of the MHC genes was linked to lower gut microbiome diversity and it was suspected that the variation in the MHC genes could affect the sensitivity to pathogens that could lead to variation in gut microbiome diversity and eventually, host survival.

To gain a better understanding of gut microbiome diversity and the contribution of different taxa to host survival, functional analyses of the gut microbiome should be included in gut microbiome studies. Different bacterial taxa can have similar functions in the gut microbiome [5, 124] and therefore, the absence of some taxa may be covered by other functionally similar taxa, resulting in a gut microbiome that is functionally more stable [125]. Similarity in functions may also contribute to host’s local adaptation e.g., to the changes in the host’s early-life environment [124]: changes in brood size or dietary items could result in variation in the gut microbiome diversity, yet there may be no effects on host body condition.

The lack of association between brood size, nestling size and survival contrasts with previous studies, but it should be noted that the majority of previous studies have been done with adult birds and not nestlings. Because nestling gut microbiome is still quite flexible compared to that of the adults [20], it is possible that our experiment did not result in a strong enough effect on the gut microbiome. In future studies, it would be important to study the parents as well as it could be more likely to find an association between adult microbiome and fitness than with nestling gut microbiome and survival. Also, our sample size in the survival analyses was small, and it is hard to determine if the result was affected by the sample size. Firstly, nestling survival is often found to correlate with brood size and more specifically, with fledging mass and in particular, the ability to forage for food [61, 126]. Intra-brood competition may explain survival to fledging, as competition between nestlings can limit food availability and thus, leading to lower nestling body condition [68, 127]. A study with blackbirds (*Turdus merula*) showed that nestling body mass explained juvenile survival [128], and similar results have been shown with great tits and collared flycatchers (*Ficedula albicollis*; [31]). Contrastingly, Ringsby et al. [129] observed that in house sparrows (*Passer domesticus*) juvenile survival was independent of nestling mass and brood size. Moreover, natal body mass is often positively correlated with survival to fledging and juvenile survival as heavier nestlings are more likely to be recruited [92, 130–131], yet we failed to demonstrate this in our study. Hatching date is also often positively correlated with fledging success [132] yet we did not find this association in our study, but instead found a significant association between hatching date and apparent juvenile survival.

## CONCLUSIONS

Offspring condition can be affected by the early-life environment and early-life gut microbiome, thus highlighting the importance of understanding how changes in the rearing environment affect individual body mass and survival. Even though our results showed between-individual variation in nestling gut microbiome diversity, we did not find a significant link between brood size and nestling gut microbiome. Moreover, we did not find a significant association between nestling gut microbiome diversity and short-term or mid-term survival. This suggests that other environmental factors (e.g., diet quality) may contribute more to variation in nestling gut microbiome. Further research is needed to uncover the environmental factors that contribute to nestling gut microbiome in wild bird populations, and how gut microbiome may be linked to nestling survival. Gut microbiome can adapt faster to environmental changes than the host, which makes it important to understand the causes of inter-individual variation in microbiome, and how variation in microbiome possibly mediate adaptation to environmental changes.

## DECLARATIONS

### Ethics statement

All animal work was conducted under relevant national and international guidelines and legislation. The animal work was licensed by the environmental and ethical committee of Varsinais-Suomi (environmental permit license number VARELY/890/2020; animal ethics permit number ESAVI/5454/2020). The birds were ringed from their nest-sites by permission from the Finnish Ringing Centre to SR (Finnish Museum of Natural History).

### Consent for publication

Not applicable

### Availability of data and material

Sequence files are available at NCBI database (BioProject PRJNA877058). Metadata and R scripts used to run the analyses are available in the Github Repository, https://github.com/marannli/bsm_analyses.

### Author contribution

The idea for this study was by AS, SR and NCS. Sample collection was done by NCS, MH, AS and SR. Laboratory analyses were done by ML with assistance from SR, KG and EV. Sequence processing and statistical analyses were done by ML with assistance from KG, EV (bioinformatics) and SR (statistics). Manuscript was written by ML and all authors commented and approved the manuscript.

## Acknowledgements

We would like to thank Charli Davies, Petri Papponen, Sari Viinikainen and Elina Virtanen at the University of Jyväskylä for their help with bioinformatics (CD) laboratory optimizations (PP, SV and EV), as well as Jorma Nurmi, Robin Cristofari, Natacha Garcin, Toni Laaksonen at the University of Turku and volunteer bird ringers for their help in the field.

## SUPPLEMENTARY INFORMATION

All supplementary files are provided in one word (.docx) document with each section titled as follows:

**Supplementary file 1.** Brood size before and after manipulation: brood sizes between treatment groups were tested with a linear model to see if the differences were statistically significant. **Supplementary file 2. Rarefaction curves for the unrarefied dataset.** Species (ASVs) plateaued at about 5000 reads which was used as the rarefying depth. **Supplementary file 3.** Phylogenetic tree using the Newick-format. The tree describes the dissimilarity among the treatment groups. Each tip represents an individual sample, and each tip is colored and shaped based on treatment. Treatment groups are clustered using the UPGMA algorithm. **Supplemental information 4.** (A) Linear mixed effects model for gut microbiome diversity (Shannon Diversity Index and Chao1 Richness) and brood size manipulation treatment. (B) Linear mixed effects model for GM diversity (Shannon Diversity Index and Chao1 Richness) and manipulated brood size**. Supplementary file 5.** A linear mixed effects model investigating the effects of brood size manipulation on nestling body mass on day 7 and day 14 post-hatch. **Supplementary file 6.** A linear mixed effects model investigating the effects of manipulated brood size on nestling body mass on day 7 and day 14 post-hatch. **Supplementary file 7.** A linear mixed effects model investigating the associations between alpha diversity (Shannon Diversity Index and Chao1 Richness) and brood size manipulation. **Supplementary file 8.** A linear mixed effects model investigating the association between alpha diversity (Shannon Diversity Index and Chao1 Richness) and manipulated brood size. **Supplementary file 9.** A generalized linear model exploration into alpha diversity’s (Shannon Diversity Index and Chao1 Richness) association with short-term (survival to fledging) and mid-term (apparent juvenile) survival. **Supplementary file 11.** Generalized linear model to measure the association between alpha diversity (Shannon Diversity Index and Chao1 Richness) survival to fledging and apparent juvenile survival. **Supplementary file 11.** Generalized linear model to measure the association between alpha diversity (Shannon Diversity Index and Chao1 Richness) survival to fledging and apparent juvenile survival. **Supplementary file 12.** Ordination of the gut microbial communities.

