## Supplementary information for "No evidence for associations between brood size, gut microbiome diversity and survival in great tit (*Parus major*) nestlings"

#### Table of Contents

Supplementary file 1. Brood size before and after manipulation. 3

Supplementary file 2. Rarefaction curves for the unrarefied dataset. 4

Supplementary file 3. Phylogenetic tree using the Newick-format. 5

Supplemental information 4. Likelihood ratio test for random effect significance 6

Supplementary file 5. A linear mixed effects model investigating the effects of brood size manipulation on nestling body mass on day 7 and day 14 post-hatch. 7

Supplemental information 5.1: Supplements for covariates and random effects 8

Supplementary file 6. A linear mixed effects model investigating the effects of final brood size on nestling body mass on day 7 and day 14 post-hatch. 9

Supplementary file 7. A linear mixed effects model investigating the associations between alpha diversity (Shannon Diversity Index and Chao1 Richness) and brood size manipulation. 10

Supplementary file 8. A linear mixed effects model investigating the association between alpha diversity (Shannon Diversity Index and Chao1 Richness) and final brood size. 11

Supplementary file 9. A generalized linear model exploration into alpha diversity's (Shannon Diversity Index and Chao1 Richness) association with short-term (survival to fledging) and mid-term (apparent juvenile) survival. 12

Supplementary file 10. The gut microbiome alpha diversity (Shannon Diversity Index and Chao1 Richness) and short-term survival. 13

Supplementary file 11. Generalized linear model to measure the association between alpha diversity (Shannon Diversity Index and Chao1 Richness) survival to fledging and apparent juvenile survival. 14

Supplementary file 12. Ordination of the gut microbial communities. 15

Supplementary file 13. Differential analysis of abundance (DESeq2) to assess the ASV abundance between the treatment groups. 16

Supplementary file 1. Brood size before and after manipulation. Brood sizes between treatment groups were tested with a linear model to see if the differences were statistically significant.

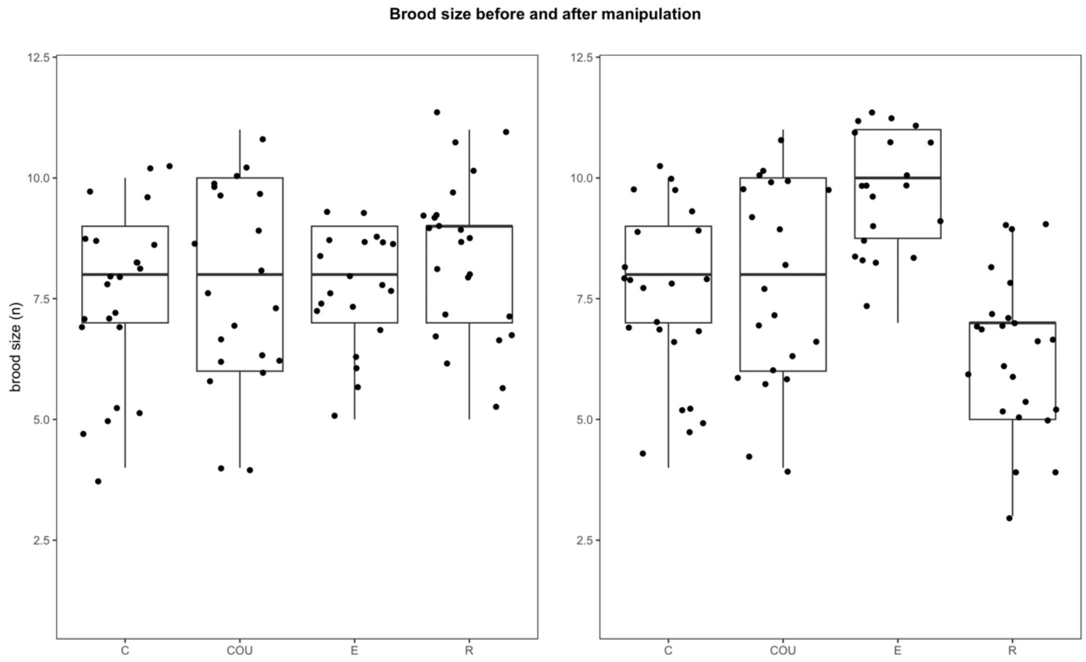

**Before brood size manipulation**

| | estimate | s.d. | *t* | p | |
|---|---|---|---|---|---|
| **(Intercept)** | 7.810 | 0.379 | 20.633 | <0.000 | *** |
| **Control treatment** | -0.244 | 0.524 | -0.467 | 0.642 | |
| **Enlarged treatment** | -0.110 | 0.542 | -0.202 | 0.840 | |
| **Reduced treatment** | 0.566 | 0.518 | 1.091 | 0.278 | |

**ANOVA**

| | df | sum.sq | mean.sq | F | p |
|---|---|---|---|---|---|
| **Treatment** | 3 | 8.910 | 2.967 | 0.987 | 0.403 |

**After brood size manipulation**

| | estimate | s.d. | *t* | p | |
|---|---|---|---|---|---|
| **(Intercept)** | 7.810 | 0.380 | 20.538 | <0.000 | |
| **Control treatment** | -0.244 | 0.526 | -0.465 | 0.643 | |
| **Enlarged treatment** | 1.841 | 0.544 | 3.380 | 0.001 | ** |
| **Reduced treatment** | -1.435 | 0.521 | -2.755 | 0.007 | ** |

**ANOVA**

| | df | sum.sq | mean.sq | F | p | |
|---|---|---|---|---|---|---|
| **Treatment** | 3 | 118.39 | 39.463 | 12.996 | <0.000 | *** |

**Tukey's post-hoc**

| **contrast** | **estimate** | **s.e.** | **df** | **t** | **p** | |
|---|---|---|---|---|---|---|
| COU-C | 0.244 | 0.526 | 84 | 0.465 | 0.967 | |
| COU-E | -1.840 | 0.544 | 84 | -3.380 | 0.006 | * |
| COU-R | 1.435 | 0.521 | 84 | 2.755 | 0.036 | * |
| C-E | -2.085 | 0.533 | 84 | -3.913 | 0.001 | ** |
| C-R | 1.190 | 0.508 | 84 | 2.341 | 0.097 | |
| E-R | 3.275 | 0.528 | 84 | 6.208 | <0.001 | *** |

Supplementary file 2. Rarefaction curves for the unrarefied dataset. Species (Amplicon Sequence Variants, ASVs) plateaued at about 5000 reads which was used as the rarefying depth.

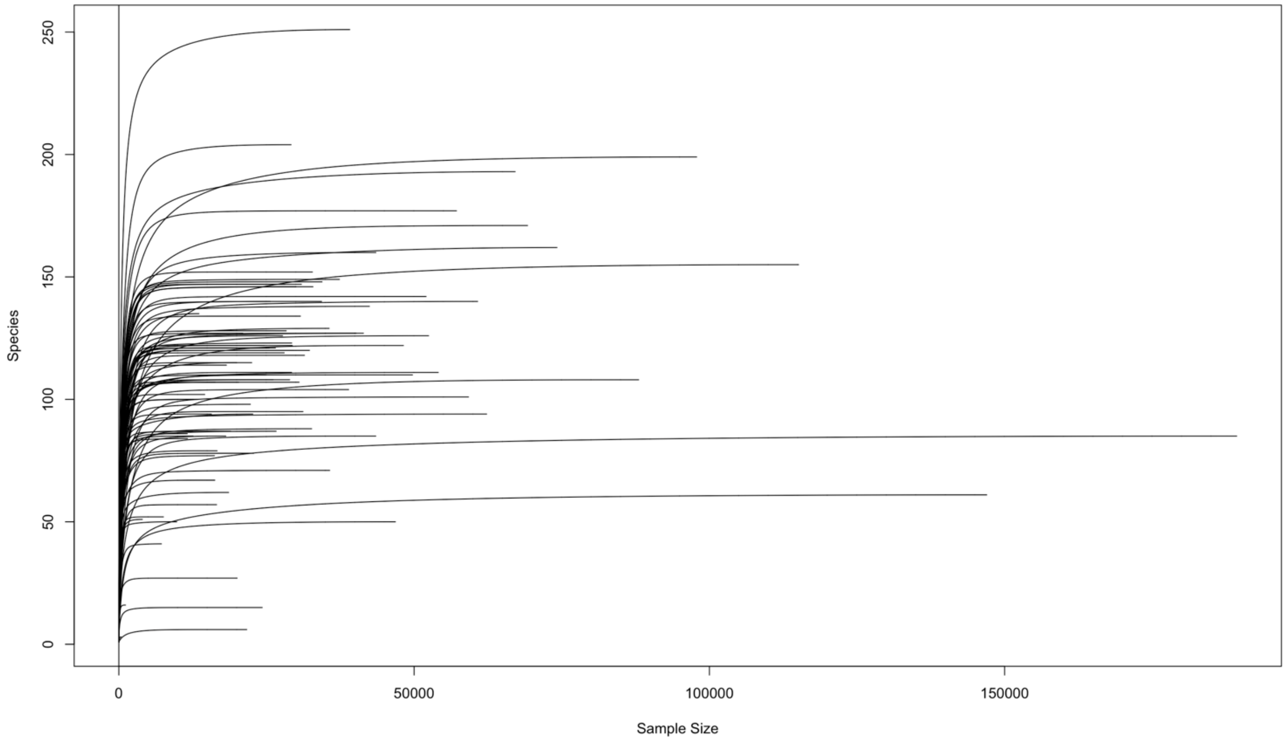

**Supplementary file 3. Phylogenetic tree using the Newick-format.** The tree describes the dissimilarity among the treatment groups. Each tip represents an individual sample, and each tip is colored and shaped based on treatment. Treatment groups are clustered using the UPGMA algorithm.

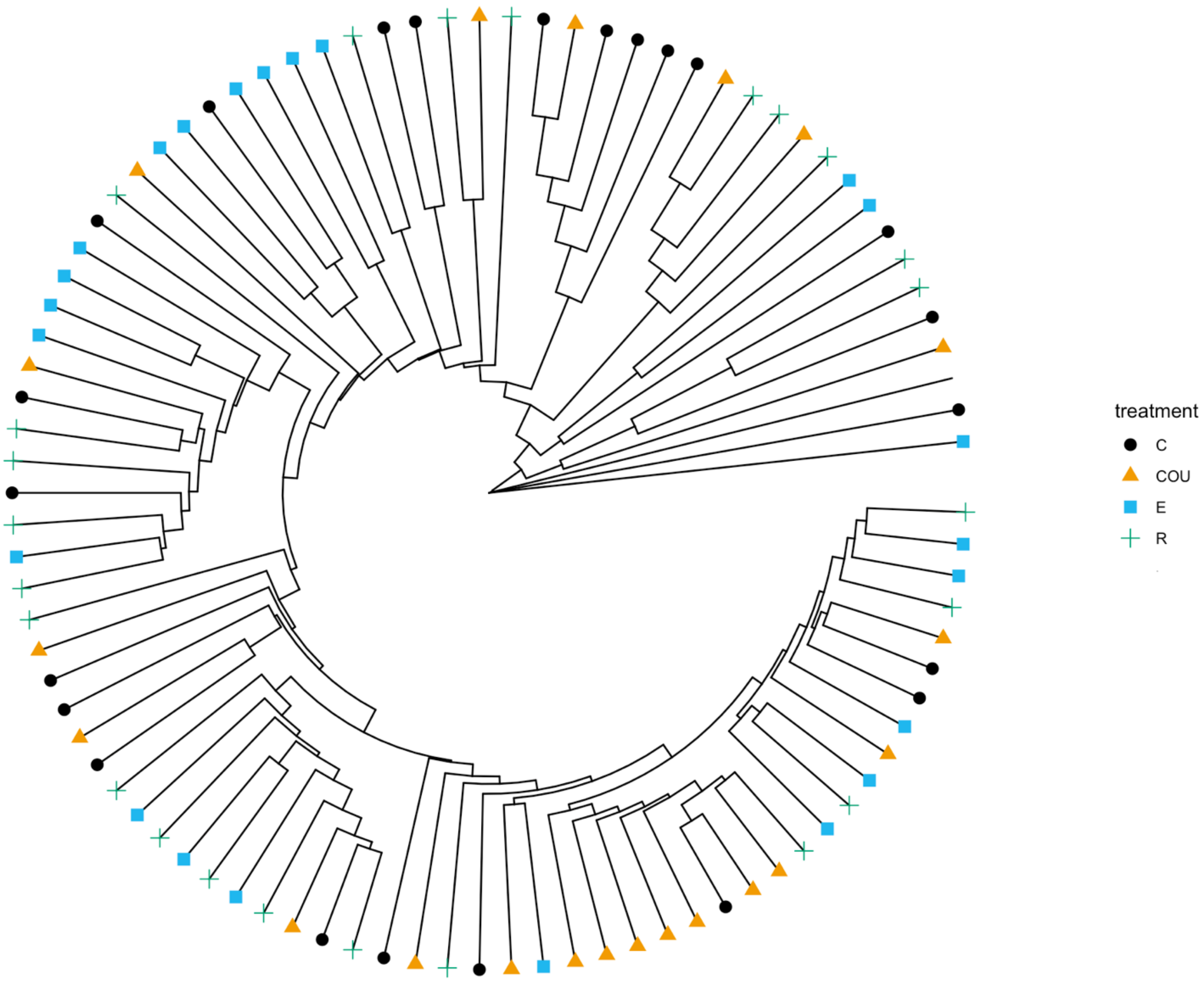

### Supplemental information 4. **Likelihood ratio test for random effect significance**

**Weight on day 7 post-hatch vs. brood size manipulation**

| Random effects | %Var | Parameters | AIC | Log lik. | Deviance | Test | $\chi2$ | Δdf | p |
|---|---|---|---|---|---|---|---|---|---|
| Nest of origin | 41.1 | 8 | 138.19 | -61.095 | 122.19 | 1 vs 3 | 0 | 1 | 1 |
| Nest of rearing | 24.4 | 8 | 138.19 | -61.095 | 122.19 | 2 vs 3 | 0 | 1 | 1 |
| Nest of origin and nest of rearing | 34.5 | 9 | 140.19 | -61.095 | 122.19 | | | | |
| Residual | | | | | | | | | |

**Weight on day 14 post-hatch vs. brood size manipulation**

| Random effects | %Var | Parameters | AIC | Log lik. | Deviance | Test | $\chi2$ | Δdf | p |
|---|---|---|---|---|---|---|---|---|---|
| Nest of origin | 65.6 | 8 | 135.17 | -59.584 | 119.17 | 1 vs 3 | 0 | 1 | 1 |
| Nest of rearing | 21.9 | 8 | 135.17 | -59.584 | 119.17 | 2 vs 3 | 0 | 1 | 1 |
| Nest of origin and nest of rearing | 12.5 | 9 | 137.17 | -59.584 | 119.17 | | | | |
| Residual | | | | | | | | | |

**Shannon Diversity Index vs brood size manipulation**

| Random effects | %Var | Parameters | AIC | Log lik. | Deviance | Test | $\chi2$ | Δdf | p |
|---|---|---|---|---|---|---|---|---|---|
| Nest of origin | 10.8 | 8 | 163.53 | -73.767 | 147.53 | 1 vs 3 | 0 | 1 | 1 |
| Nest of rearing | 27.7 | 8 | 163.53 | -73.767 | 147.53 | 2 vs 3 | 0 | 1 | 1 |
| Nest of origin and nest of rearing | 31.1 | 9 | 165.53 | -73.767 | 147.53 | | | | |
| Residual | | | | | | | | | |

**Chao1 Richness vs brood size manipulation**

| Random effects | %Var | Parameters | AIC | Log lik. | Deviance | Test | $\chi2$ | Δdf | p |
|---|---|---|---|---|---|---|---|---|---|
| Nest of origin | 10.7 | 8 | 698.65 | -341.32 | 682.65 | 1 vs 3 | 0 | 1 | 1 |
| Nest of rearing | 23.9 | 8 | 698.65 | -341.32 | 682.65 | 2 vs 3 | 0 | 1 | 1 |
| Nest of origin and nest of rearing | 65.4 | 9 | 700.65 | -341.32 | 682.65 | | | | |
| Residual | | | | | | | | | |

**Supplementary file 5.** A linear mixed effects model investigating the effects of brood size manipulation on nestling body mass on day 7 and day 14 post-hatch.

| **Weight D7** | | | | | | |
|---|---|---|---|---|---|---|
| | **estimate** | **s.e.** | **df** | ***t*** | **p** | |
| **(Intercept)** | 5.93456 | 2.582 | 25.522 | 2.298 | 0.030 | * |
| **Enlarged brood size** | -0.387 | 0.595 | 26.154 | -0.649 | 0.522 | |
| **Reduced brood size** | 0.146 | 0.518 | 24.937 | 0.282 | 0.780 | |
| **Original brood size** | -0.075 | 0.159 | 26.391 | -0.473 | 0.640 | |
| **Hatching date** | 0.057 | 0.039 | 24.444 | 1.436 | 0.164 | |
| **Weight D2** | 0.662 | 0.146 | 31.682 | 4.522 | < 0.000 | *** |
| **(Interactions)** | | | | | | |
| **(Enlarged * original brood size** | -0.105 | 0.421 | 25.254 | -0.251 | 0.804) | |
| **(Reduced * original brood size** | 0.322 | 0.370 | 23.902 | 0.871 | 0.393) | |

| **Random effects** | | |
|---|---|---|
| | **variance** | **s.d.** |
| **Nest of origin** | 0.668 | 0.817 |
| **Nest of rearing** | 0.396 | 0.629 |
| **Residual** | 0.560 | 0.749 |

| **Weight D14** | | | | | | |
|---|---|---|---|---|---|---|
| | **estimate** | **s.e.** | **df** | ***t*** | **p** | |
| **(Intercept)** | 5.241 | 3.290 | 24.404 | 1.593 | 0.124 | |
| **Enlarged brood size** | -0.527 | 0.778 | 24.655 | -0.678 | 0.504 | |
| **Reduced brood size** | 0.359 | 0.684 | 23.788 | 0.526 | 0.604 | |
| **Original brood size** | 0.093 | 0.194 | 24.460 | 0.479 | 0.636 | |
| **Hatching date** | 0.184 | 0.050 | 24.070 | 3.656 | 0.001 | ** |
| **Weight D2** | 0.089 | 0.167 | 30.925 | 0.532 | 0.599 | |
| **(Interactions)** | | | | | | |
| **(Enlarged * original brood size** | 0.054 | 0.521 | 22.643 | 0.104 | 0.918) | |
| **(Reduced * original brood size** | 0.316 | 0.464 | 22.215 | 0.681 | 0.503) | |

| **Random effects** | | |
|---|---|---|
| | **variance** | **s.d.** |
| **Nest of origin** | 1.525 | 1.235 |
| **Nest of rearing** | 0.510 | 0.714 |
| **Residual** | 0.291 | 0.539 |

This basic model includes control (C), enlarged (E), and reduced (R) groups. Interactions between manipulated brood size and original brood size were removed from final models as there was no significant interaction and are shown in the table below. Nest of origin and nest of rearing were included as random effects to control for the non-independency of samples.

#### Supplemental information 5.1: Supplements for covariates and random effects

*The effects of brood size manipulation on nestling body mass*

Original brood size (ANOVA: $F_{1, 27.486}=0.189$, $p=0.667$, Table 1) and hatching date (ANOVA: $F_{1, 25.813}=1.712$, $p=0.202$, Table S1) did not associate with nestling body mass on day 7 post-hatch. Weight on day 2 positively correlated with weight on day 7 post hatch (ANOVA: $F_{1, 10.884}=33.697$, $p=<0.000$, Table S1). Variance explained by the nest of origin was estimated to be higher ($\sigma2=0.698$, s.d.=0.836) than by the nest of rearing ($\sigma2=0.417$, s.d.=0.646). Weight on day 2 post-hatch did not affect nestling body weight on day 14 post-hatch (ANOVA: $F_{1, 10.600}=0.032$, $p=0.861$, Table S1). However, hatching date was positively correlated with nestling body weight on day 14 post-hatch (ANOVA: $F_{1, 24.345}=11.903$, $p=0.002$, Table 1). There was no significant interaction between brood size manipulation and original brood size (ANOVA: $F_{2, 23.223}=0.219$, $p=0.805$, Table S1). Variance explained by the nest of origin was lower ($\sigma2=0.865$, s.d.=0.930) than by the nest of rearing ($\sigma2=0.996$, s.d.=0.840).

**Supplementary file 6.** A linear mixed effects model investigating the effects of final brood size on nestling body mass on day 7 and day 14 post-hatch.

| **Weight D7** | | | | | | |
|---|---|---|---|---|---|---|
| | **estimate** | **s.e.** | **df** | ***t*** | **p** | |
| **(Intercept)** | 6.383 | 2.488 | 33.280 | 2.566 | 0.015 | * |
| **Manipulated broodsize** | -0.151 | 0.113 | 35.149 | -1.333 | 0.191 | |
| **Hatching date** | 0.053 | 0.038 | 32.724 | 1.386 | 0.175 | |
| **Weight D2** | 0.717 | 0.149 | 40.007 | 4.808 | <0.000 | *** |
| **(Interactions)** | | | | | | |
| **(Manipulated brood size * Hatching date** | -0.027 | 0.025 | 35.140 | -1.094 | 0.281) | |

| **Random effects** | | |
|---|---|---|
| | **variance** | **s.d.** |
| **Nest of origin** | 1.332 | 1.154 |
| **Residual** | 0.561 | 0.749 |

| **Weight D14** | | | | | | |
|---|---|---|---|---|---|---|
| | **estimate** | **s.e.** | **df** | ***t*** | **p** | |
| **(Intercept)** | 6.671 | 3.072 | 28.882 | 2.171 | 0.038 | * |
| **Manipulated broodsize** | -0.100 | 0.125 | 29.641 | -0.796 | 0.432 | |
| **Hatching date** | 0.183 | 0.048 | 28.278 | 3.804 | 0.001 | ** |
| **Weight D2** | 0.100 | 0.155 | 36.871 | 0.643 | 0.524 | |
| **(Interactions)** | | | | | | |
| **(Manipulated brood size * Hatching date** | -0.017 | 0.028 | 32.189 | -0.613 | 0.544) | |

| **Random effects** | | |
|---|---|---|
| | **variance** | **s.d.** |
| **Nest of origin** | 1.999 | 1.414 |
| **Residual** | 0.286 | 0.535 |

The analysis includes all treatment groups i.e., the full model: control (C), unmanipulated control (COU), enlarged (E), and reduced (R). Nest of origin was included as a random effect to control for the non-independency of samples.

**Supplementary file 7.** A linear mixed effects model investigating the associations between alpha diversity (Shannon Diversity Index and Chao1 Richness) and brood size manipulation.

| Shannon Diversity Index | | | | | |
|---|---|---|---|---|---|
| | **estimate** | **s.e.** | **df** | ***t*** | **p** |
| **(Intercept)** | 1.572 | 1.201 | 51.928 | 1.309 | 0.196 |
| **Control brood size** | 0.345 | 0.285 | 42.108 | 1.213 | 0.232 |
| **Enlarged brood size** | 0.495 | 0.286 | 48.364 | 1.733 | 0.090 |
| **Reduced brood size** | 0.291 | 0.274 | 46.894 | 1.061 | 0.294 |
| **Original brood size** | -0.035 | 0.056 | 50.269 | -0.623 | 0.536 |
| **Weight D7** | -0.003 | 0.051 | 80.551 | -0.052 | 0.959 |
| **Hatching date** | 0.018 | 0.018 | 50.276 | 1.036 | 0.305 |
| **(Interactions)** | | | | | |
| **(Control treatment * Original brood size** | -0.070 | 0.142 | 44.595 | -0.489 | 0.627) |
| **(Enlarged treatment * Original brood size** | -0.067 | 0.181 | 53.253 | -0.370 | 0.713) |
| **(Reduced treatment * Original brood size** | 0.005 | 0.152 | 44.231 | 0.030 | 0.976) |

| Random effects | | |
|---|---|---|
| | **Variance** | **s.d.** |
| **Nest of rearing** | 0.229 | 0.479 |
| **Residual** | 0.388 | 0.623 |

| Chao1 Richness | | | | | |
|---|---|---|---|---|---|
| | **estimate** | **s.e.** | **df** | ***t*** | **p** |
| **(Intercept)** | 94.200 | 60.218 | 49.224 | 1.564 | 0.124 |
| **Control brood size** | 6.219 | 13.979 | 38.526 | 0.445 | 0.659 |
| **Enlarged brood size** | -5.698 | 14.238 | 46.609 | -0.400 | 0.691 |
| **Reduced brood size** | -5.014 | 13.621 | 44.559 | -0.368 | 0.715 |
| **Original brood size** | -1.541 | 2.811 | 50.403 | -0.548 | 0.586 |
| **Weight D7** | -2.127 | 2.746 | 80.488 | -0.775 | 0.441 |
| **Hatching date** | 0.897 | 0.878 | 49.214 | 1.022 | 0.312 |
| **(Interactions)** | | | | | |
| **(Control treatment * Original brood size** | 5.707 | 7.091 | 45.209 | 0.805 | 0.425) |
| **(Enlarged treatment * Original brood size** | 6.394 | 8.186 | 56.608 | 0.696 | 0.489) |
| **(Reduced treatment * Original brood size** | 4.995 | 7.566 | 44.066 | 0.660 | 0.513) |

| Random effects | | |
|---|---|---|
| | **Variance** | **s.d.** |
| **Nest of rearing** | 269.9 | 16.42 |
| **Residual** | 1424.4 | 37.74 |

The model includes all four treatment groups i.e., the full model: control (C), unmanipulated control (COU), enlarged (E), and reduced (R). Interactions between brood size manipulation and original brood size were removed as there was no significant interaction and are shown in the table below. Nest of rearing was included as a random effect to control for the non-independency of samples.

**Supplementary file 8.** A linear mixed effects model investigating the association between alpha diversity (Shannon Diversity Index and Chao1 Richness) and final brood size.

**Shannon Diversity Index**

| | estimate | s.e. | df | *t* | p | |
|---|---|---|---|---|---|---|
| **(Intercept)** | 2.349 | 1.053 | 65.137 | 2.231 | 0.029 | * |
| **Manipulated brood size** | <0.000 | 0.045 | 63.001 | 0.020 | 0.984 | |
| **Weight D7** | -0.006 | 0.051 | 82.840 | -0.123 | 0.903 | |
| **Hatching date** | 0.006 | 0.016 | 59.734 | 0.370 | 0.713 | |
| **(Interactions)** | | | | | | |
| **(Manipulated brood size* weight D7** | -0.001 | 0.022 | 82.270 | -0.003 | 0.998) | |

**Random effects**

| | Variance | s.d. |
|---|---|---|
| **Nest of rearing** | 0.248 | 0.498 |
| **Residual** | 0.377 | 0.614 |

**Chao1 Richness**

| | estimate | s.e. | df | *t* | p |
|---|---|---|---|---|---|
| **(Intercept)** | 93.862 | 52.199 | 64.008 | 1.791 | 0.078 |
| **Manipulated brood size** | -1.099 | 2.215 | 65.064 | -0.496 | 0.622 |
| **Weight D7** | -2.249 | 2.709 | 83.513 | -0.830 | 0.409 |
| **Hatching date** | 0.849 | 0.798 | 57.110 | 1.064 | 0.292 |
| **(Interactions)** | | | | | |
| **(Manipulated brood size * weight D7** | -0.683 | 1.176 | 81.896 | -0.580 | 0.563) |

**Random effects**

| | Variance | s.d. |
|---|---|---|
| **Nest of rearing** | 238.7 | 15.45 |
| **Residual** | 1419.0 | 37.67 |

The model includes all four treatment groups i.e., the full model: control (C), unmanipulated control (COU), enlarged (E), and reduced (R). Interactions between alpha diversity and final brood size were removed as there was no significant interaction and are shown in the table below. Nest of rearing was included as a random effect to control for the non-independency of samples.

**Supplementary file 9.** A generalized linear model exploration into alpha diversity's (Shannon Diversity Index and Chao1 Richness) association with short-term (survival to fledging) and mid-term (apparent juvenile) survival.

**Survival to fledging (Shannon Diversity Index)**

| | estimate | s.e. | z value | p |
|---|---|---|---|---|
| **(Intercept)** | -1.505 | 4.163 | -0.362 | 0.718 |
| **Shannon** | -0.044 | 0.457 | -0.097 | 0.923 |
| **Weight D7** | 0.143 | 0.230 | 0.622 | 0.765 |
| **Hatching date** | 0.043 | 0.064 | 0.669 | 0.503 |
| **Manipulated brood size** | -0.057 | 0.190 | -0.298 | 0.765 |

**Survival to fledging (Chao1 Richness)**

| | estimate | s.e. | z value | p |
|---|---|---|---|---|
| **(Intercept)** | -1.312 | 4.215 | -0.311 | 0.756 |
| **Chao1** | -0.003 | 0.009 | -0.343 | 0.732 |
| **Weight D7** | 0.134 | 0.230 | 0.584 | 0.559 |
| **Hatching date** | 0.046 | 0.065 | 0.706 | 0.480 |
| **Manipulated brood size** | -0.062 | 0.192 | -0.321 | 0.748 |

**Recapture as juvenile (Shannon Diversity Index)**

| | estimate | s.e. | z value | p | |
|---|---|---|---|---|---|
| **(Intercept)** | 2.780 | 3.388 | 0.821 | 0.412 | |
| **Shannon** | -0.501 | 0.368 | -1.362 | 0.173 | |
| **Weight D7** | 0.270 | 0.185 | 1.460 | 0.144 | |
| **Hatching date** | -0.102 | 0.049 | -2.087 | 0.037 | * |
| **Manipulated brood size** | 0.014 | 0.143 | 0.096 | 0.923 | |

**Recapture as juvenile (Chao1 Richness)**

| | estimate | s.e. | z value | p | |
|---|---|---|---|---|---|
| **(Intercept)** | 4.214 | 3.409 | 1.236 | 0.216 | |
| **Chao1** | -0.013 | 0.008 | -1.559 | 0.119 | |
| **Weight D7** | 0.204 | 0.199 | 1.023 | 0.306 | |
| **Hatching date** | -0.109 | 0.052 | -2.104 | 0.035 | * |
| **Manipulated brood size** | 0.023 | 0.146 | 0.161 | 0.872 | |

*Random effects were excluded as the model failed to converge.

**Supplementary file 10.** The gut microbiome alpha diversity (Shannon Diversity Index and Chao1 Richness) and short-term survival.

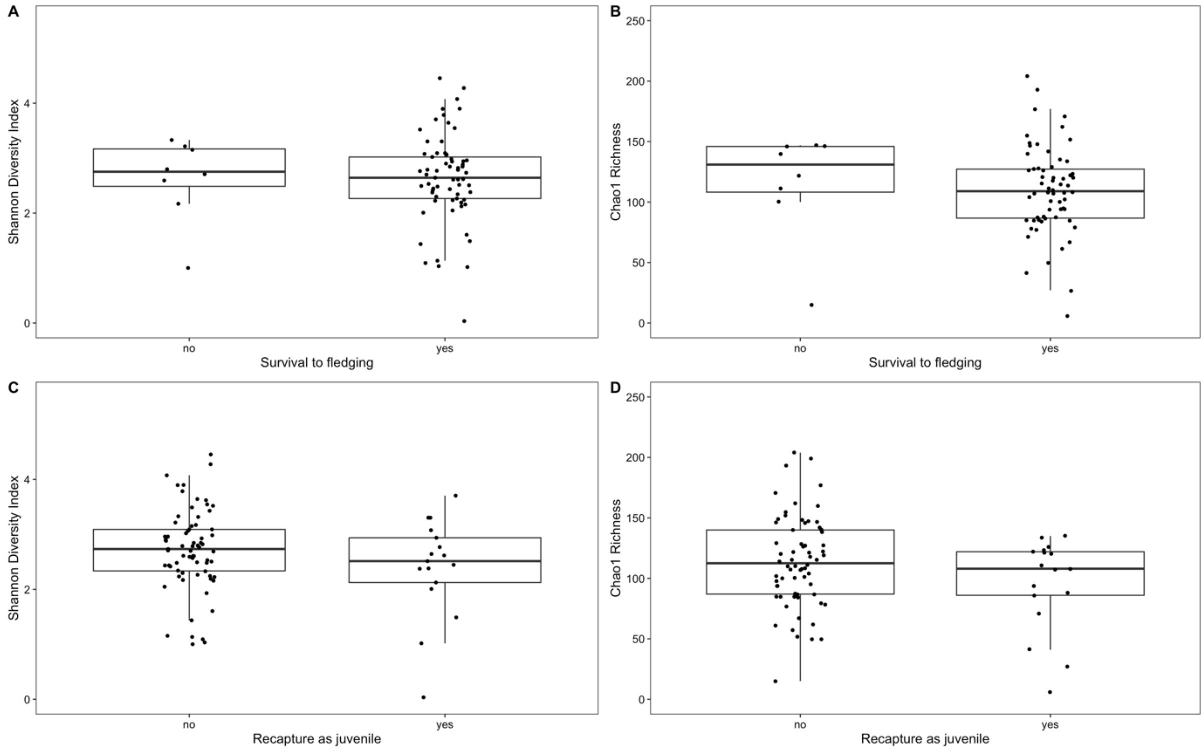

In survival to fledging (A: Shannon Diversity Index; B: Chao1 Richness) 65 nestlings fledged successfully and 8 nestlings were dead. 15 nestlings had no fledging record, so these were excluded from the analysis. In recapture as juvenile (C: Shannon Diversity Index; D: Chao1 Richness) 19 out of 92 (with data on microbiome diversity) were captured. The black dots represent each observation within a treatment group. The whiskers represent 95 % confidence intervals.

**Supplementary file 11.** Generalized linear model to measure the association between alpha diversity (Shannon Diversity Index and Chao1 Richness) survival to fledging and apparent juvenile survival.

**Survival to fledging without the unmanipulated control group (COU)**

| | estimate | s.e. | z value | p |
|---|---|---|---|---|
| **(Intercept)** | -3.901 | 7.450 | -0.524 | 0.601 |
| **Shannon** | -1.153 | 0.889 | -1.297 | 0.195 |
| **Weight D7** | 0.389 | 0.469 | 0.830 | 0.407 |
| **Manipulated brood size** | -0.624 | 0.410 | -1.523 | 0.128 |
| **Hatching date** | 0.185 | 0.112 | 1.648 | 0.100 |

| | estimate | s.e. | z value | p |
|---|---|---|---|---|
| **(Intercept)** | -3.718 | 6.265 | -0.593 | 0.553 |
| **Chao1** | -0.011 | 0.013 | -0.809 | 0.419 |
| **Weight D7** | 0.255 | 0.385 | 0.664 | 0.507 |
| **Manipulated brood size** | -0.557 | 0.388 | -1.437 | 0.151 |
| **Hatching date** | 0.161 | 0.107 | 1.507 | 0.132 |

**Recapture as juvenile without nestlings that had no survival to fledging datapoint**

| | estimate | s.e. | z value | p | |
|---|---|---|---|---|---|
| **(Intercept)** | 3.816 | 3.355 | 1.137 | 0.256 | |
| **Shannon** | -0.455 | 0.378 | -1.203 | 0.229 | |
| **Weight D7** | 0.241 | 0.192 | 1.254 | 0.210 | |
| **Manipulated brood size** | 0.043 | 0.146 | 0.293 | 0.770 | |
| **Hatching date** | -0.116 | 0.051 | -2.275 | 0.023 | * |

| | estimate | s.e. | z value | p | |
|---|---|---|---|---|---|
| **(Intercept)** | 4.214 | 3.409 | 1.236 | 0.216 | |
| **Chao1** | -0.013 | 0.008 | -1.559 | 0.119 | |
| **Weight D7** | 0.204 | 0.199 | 1.023 | 0.306 | |
| **Manipulated brood size** | 0.023 | 0.146 | 0.161 | 0.872 | |
| **Hatching date** | -0.109 | 0.052 | -2.104 | 0.035 | * |

The survival to fledging model is a basic model including the control (C), enlarged (E), and reduced (R) treatment groups and individuals with no recorded fledging success removed. In the recapture as juvenile model, nestlings without a recorded fledging success (N=16) are removed from the data. To simplify both models, all random intercepts (nest of origin and nest of rearing) were removed as they contained 59 levels each and the model failed to converge when they were included.

**Supplementary file 12.** Ordination of the gut microbial communities.

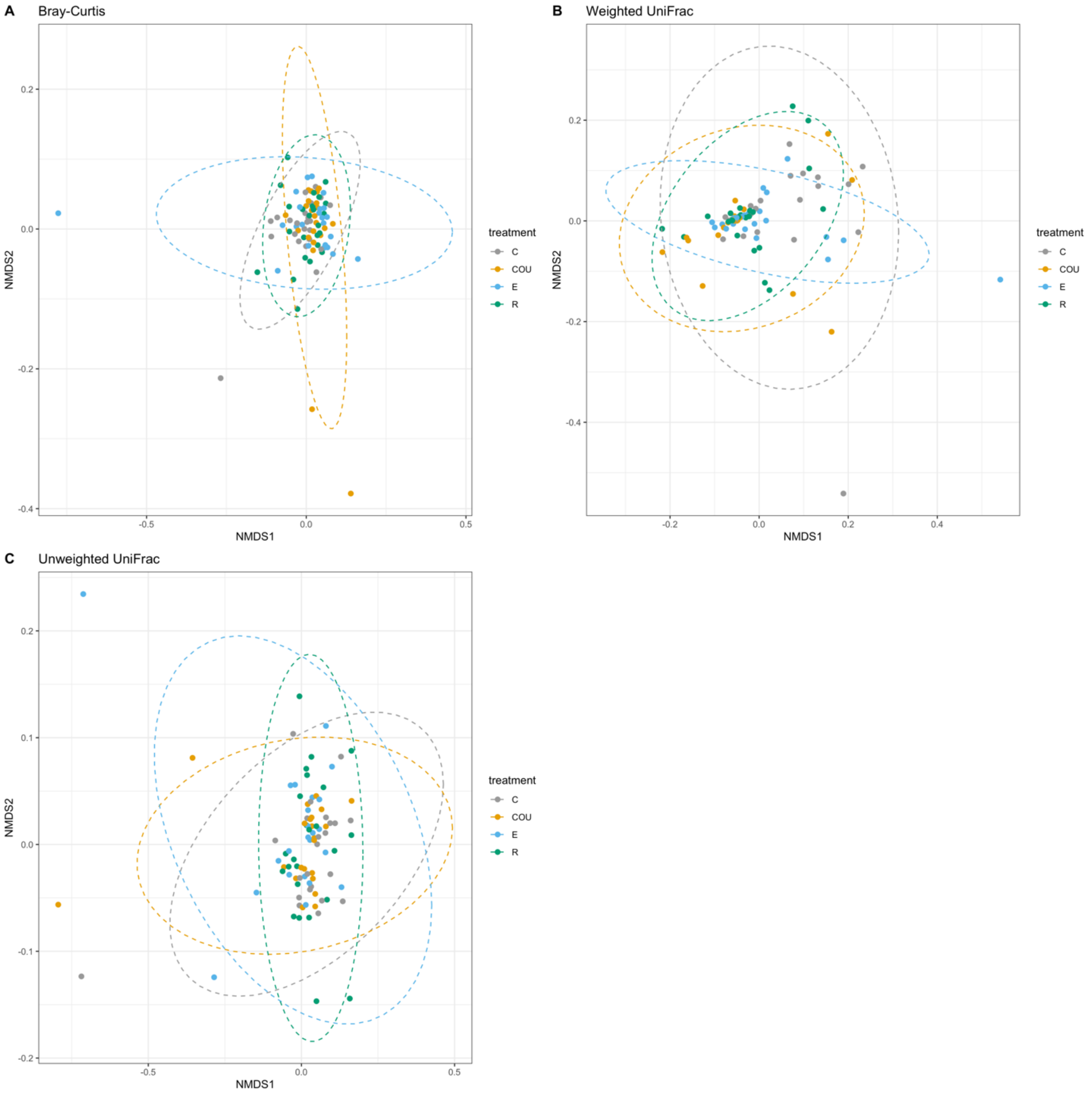

A) Weighted UniFrac, B) Unweighted UniFrac, and C) Bray-Curtis dissimilarity are displayed on NMDS ordinations. The color of the dots indicates which treatment, and the dashed ellipses represent 95 % confidence intervals.

#### Supplementary file 13. Differential analysis of abundance (DESeq2) to assess the ASV abundance between the treatment groups

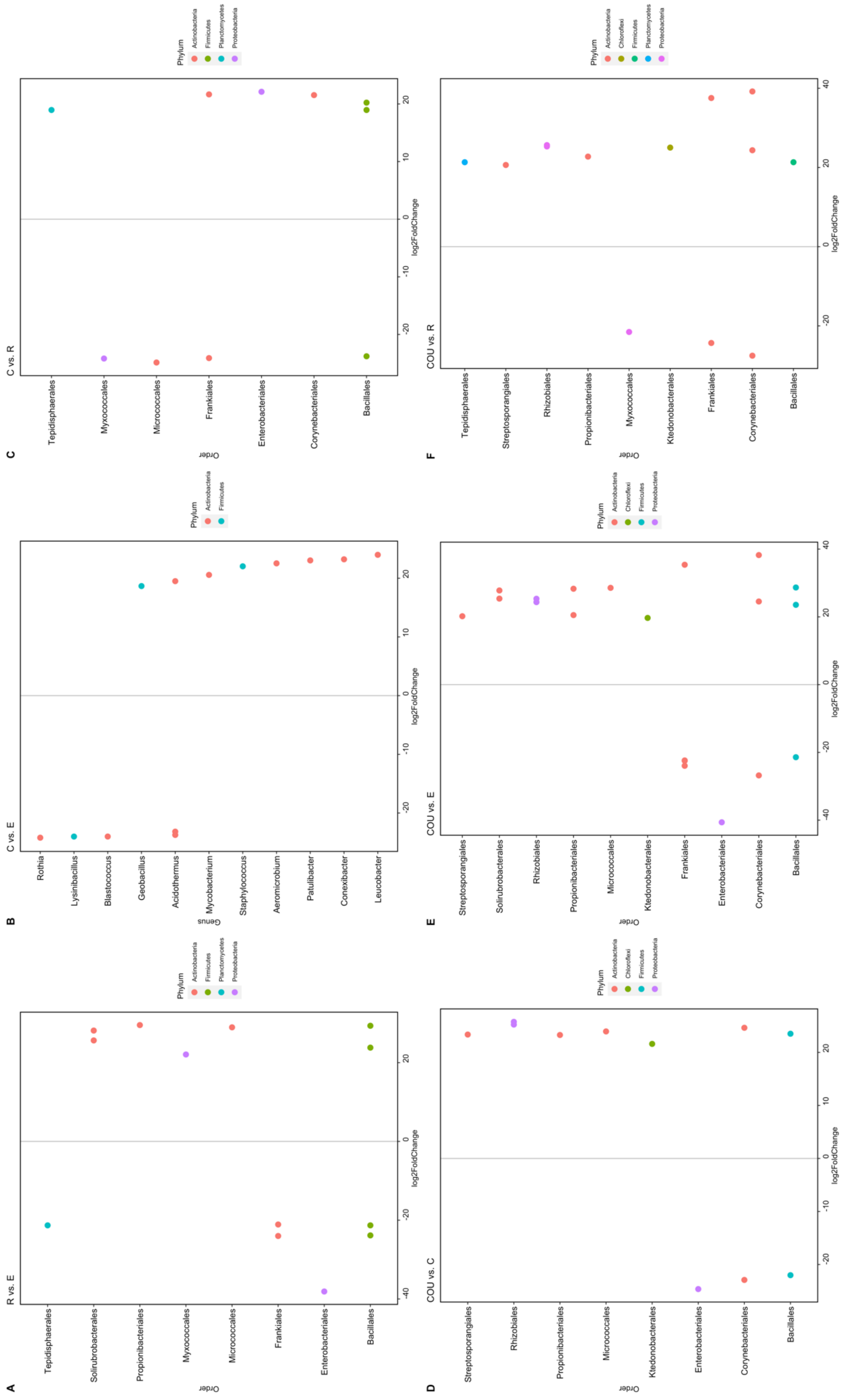
